# Sex differences in pant-hoot vocalizations in wild Eastern chimpanzees

**DOI:** 10.1101/2024.12.14.628485

**Authors:** Eve Holden, Charlotte Grund, Robert Eguma, Liran Samuni, Klaus Zuberbühler, Adrian Soldati, Catherine Hobaiter

## Abstract

Chimpanzee pant-hoots are frequently produced long-distance vocalisations that serve a number of social functions, such as to indicate coordinate travelling, food discovery, or social status. Calling often triggers replies by others, offering potential for some form of group-level decision-making through vocal exchanges. However, despite important physiological and social differences between male and female chimpanzees, pant-hoot research has traditionally focused on males, with very little known about female use. To address this gap, we collected all-occurrence behavioural data from wild adult female and male Eastern chimpanzees (*Pan troglodytes schweinfurthii*) in two communities in the Budongo Forest Reserve, Uganda. We show that females pant-hooted regularly, but less frequently than males and, when they did so, mostly in response to others. Response pant-hoots were more common than initiating pant-hoots in both sexes (females: 90.1%; males: 69.5%), and male and female chimpanzees responded to pant-hoots heard at a similar rate (females: 14.1%, males excluding alpha: 15.8%). Both sexes were more likely to respond to pant-hoots from within their own party, but this effect was stronger for females. The likelihood of male chimpanzees pant-hooting in response was inhibited in larger party sizes, while females’ response pant-hoots were not. Finally, female response pant-hoots were less common during periods of maximal oestrous as compared to other stages of their reproductive cycle. Overall, our findings demonstrate that response pant-hoots are subject to social factors, such as within-sex social competition and between-sex reproductive strategies, which affect male and female chimpanzees in different ways, a further demonstration of the high degree of audience awareness in this primate species.

**Highlights:**

- Chimpanzees use pant-hoots in both sex-specific and sex-non-specific ways
- Individual and social factors impact likelihood of pant-hoot production
- Our findings suggest pant-hoots are important flexible elements of vocal behaviour for male and female chimpanzees.

## Introduction

Animals encode varied information when using vocal signals to communicate with each other (Bradbury & Vehrencamp, 2011). By observing and quantifying patterns of behavioural response and the socio-ecological factors associated with signal production, researchers can attempt to better understand signal function. For example, diverse taxa use vocalisations in ways that inform others about threats (e.g., Hollén & Radford, 2009) and food availability (e.g., Clay et al., 2012), and the acoustic structure of some calls encodes specific information about the caller’s identity (e.g., Aubin et al., 2000), and/or their activity (e.g., McComb & Reby, 2009). When groups are dispersed across large territories and communication takes place over longer distances, having structurally distinct calls to maintain auditory contact and coordinate movements with distant conspecifics can be particularly beneficial—especially when visual contact is difficult due to dense or low-visibility habitats (e.g., Amazonian manatees, *Trichechus inunguis*: Sousa-Lima et al., 2002; elephants, *Loxodonta africana*: McComb et al., 2003; howler monkeys, *Alouatta palliata*: Ceccarelli et al., 2021; spotted hyenas, *Crocuta Crocuta*: Gersick et al., 2015). In species where social dynamics are characterised by a high degree of fission-fusion dynamics (i.e., where group members split from and reunite with others regularly throughout the day) individuals may benefit by using and responding to long-distance calls. The fact that these calls can encode features such as the identity, location, and activity of the caller (Tibbetts & Dale, 2007)—could allow individuals to make informed decisions about where and when to travel, who to associate with, and how to adjust their social decisions depending on who is involved, for instance during disputes over territory or status (e.g., bonobos, *Pan paniscus*: White et al., 2015; elephants, *Loxodonta africana:* Leighty et al., 2008; lions, *Pantera leo*: McComb et al., 1994). It is also adaptive for individuals in more cohesive social units or contexts to consider the socio-ecological correlates of a communicative event in order to anticipate external events (e.g., predators or other threats), or to reduce uncertainty about other individuals’ status and behaviour (Seyfarth et al., 2010). Hamadryas baboons (*Papio hamadryas ursinus*) keep track of dominance hierarchies by inferring the role of a caller during an agonistic interaction (Bergman et al., 2003), and bottlenose dolphins (*Tursiops truncates*) comfort or reconcile with conspecifics in response to hearing their distress calls (Kuczaj et al., 2015).

Some animals adjust their vocal output based on the presence or absence of specific individuals in their vicinity or immediate social group, so-called ‘audience effects’ (e.g., chickens, *Gallus gallus*: Evans & Marler, 1994; chimpanzees, *Pan troglodytes*: Crockford et al., 2017; Siamese fighting fish, *Betta splendens*: Matos et al., 2003; tufted capuchin monkeys, *Cebus apella nigritus*: Di Bitetti, 2005; yellow mongoose, *Cynictis penicillata*: Le Roux et al., 2008; zebra finches, *Taeniopygia guttata* Vignal et al., 2004). Audience effects, which may be shaped by the presence of both nearby and distant receivers (Evans, 1997), demonstrate that signallers have a level of flexibility in their production (or inhibition) of vocalizations (Zuberbühler, 2008). Primates’ rich multi-layered social interactions and relationships vary greatly within and across species and are primary drivers of individual survival and success (Cheney & Seyfarth, 2018; Silk, 2007). There is growing evidence that an increasing number of primate species demonstrate flexible control of their vocal production in ways that might help to mediate social relationships (Freeberg et al., 2012; Gustison et al., 2012). Flexibility in vocal production can be measured at the level of spontaneous use, i.e., whether signallers call depending on socio-ecological context or at the level of response, i.e., whether receivers call in response to the calls of others (Seyfarth & Cheney, 2003, 2018).

Highly gregarious, chimpanzees live in multi-male multi-female social groups with fission-fusion dynamics (Aureli et al., 2008), sometimes blended with additional levels of social organisation where group sizes are atypically large (e.g., Ngogo: Mitani & Amsler, 2003; Waibira: Badihi et al., 2022). In the wild, chimpanzees occupy large territories in which individuals associate in short-term ‘parties’ that change in size and composition across the day (Nishida, 1968; Van Lawick-Goodall, 1968). To aid in navigating where and when to associate with individuals in their community across large and often densely-forested areas, chimpanzees employ a long-distance vocalisation: the pant-hoot, from which they can accurately track the location and identity of the caller at over a kilometre away (Ghiglieri, 2019; Reynolds, 2005; Wilson et al., 2001). Pant-hoots consist of up to four distinct phases, defined by the acoustic characteristics of the elements within them (Marler & Hobbett, 1975): introduction, build-up, climax, and let-down. Different acoustic elements within these phases encode callers’ identity, age, social status, and activity (Crockford et al., 2004; Desai et al., 2022; Fedurek et al., 2016; Kojima et al., 2003), providing a useful mechanism to recruit party members and coordinate movements across distant parties. For example, once reaching a plentiful or high-quality feeding patch, male chimpanzees often produce an acoustically distinct pant-hoot let-down phase (Fedurek et al., 2016; Notman & Rendall, 2005). Pant-hoots are also frequently combined with ‘drumming’ on the large buttresses of trees (Arcadi et al., 1998), which encodes further information on identity, activity, and location (Eleuteri et al., 2022). In experimental studies, patterns of pant-hoot responses during simulated inter-community encounters suggested that chimpanzees use these calls to assess group strength and whether to engage, depending on their own group size and composition (Herbinger et al., 2009; Wilson et al., 2001).

In addition to variation in their acoustic structure, the production of pant-hoots appears to be sensitive to audience composition, early life exposure, and other social factors (Bründl et al., 2022; Fedurek, Machanda, et al., 2013; Mitani & Nishida, 1993; Soldati et al., 2022). Because chimpanzees show flexibility in their production of and responses to pant-hoots across behavioural contexts, these calls offer a basis from which to investigate the interplay of social and environmental effects on vocal usage, including the impact of social and individual differences in calling (e.g. Soldati et al., under review), and sex-specific variation. At present, despite over 50-years of interest (Marler & Hobbett, 1975) and a substantial literature, almost all pant-hoot studies focus exclusively on their use by adult males. As in a range of other animal communication systems (e.g., songbirds: Odom et al., 2014), female chimpanzees represent the ‘forgotten sex’ in the study of vocalizations, with just a handful of studies on their short-to-medium distance copulation or greeting calls (Fallon et al., 2016; Laporte & Zuberbühler, 2010; Townsend et al., 2008, 2011). One reason for this bias in studies of pant-hoots is that early work suggested (from very small samples) that female chimpanzees rarely produced pant-hoots and lacked the characteristic climax phase (Marler & Hobbett, 1975), regarded by many as a defining acoustic feature (e.g., Arcadi, 1996; Fedurek et al., 2014; Mitani et al., 1999; Mitani & Brandt, 1994). Recently this definition has become untenable: mature male chimpanzees also regularly produce pant-hoots that lack a climax phase (Soldati et al., 2022) and females have been observed producing pant-hoots with and without climax phases (Crunchant et al., 2021; Kalan, 2019; Wilson et al., 2007). Another factor in their less frequent study, is the difficulty of collecting data: female chimpanzees are generally less vocal than males and, when they produce pant-hoots, it is often as part of a pant-hoot response in which multiple individual’s calls are produced in ‘chorus’ together (Clark, 1993; Marler & Tenaza, 1977; Notman & Rendall, 2005), making acoustic analysis challenging. Moreover, as compared to males, the longer habituation period needed to observe some females (sometimes over a decade: Boesch-Achermann & Boesch, 1994) and their more solitary lifestyle in Eastern chimpanzees (*Pan troglodytes schweinfurthii*; Gilby & Wrangham, 2008; Lehmann & Boesch, 2008) necessitate substantial investment in data collection. In sum, while there is a growing consensus that adult female chimpanzees produce pant-hoots, there has also been no systematic study of their use of this important vocal signal.

Studying female vocal behaviour does more than provide a more informed picture of chimpanzee vocal behaviour as a whole. In many primate species, the use of vocalisations is shaped by sex-specific aspects of physiology and behaviour—making it inappropriate to make species generalisations from the behaviour of one of the sexes. For example, female yellow baboons (*Papio cynocephalus cynocephalus*) encode their oestrus state in copulation calls (Semple et al., 2002), and rhesus macaques (*Macaca mulatta*) alter vocal responses depending on the sex of the conspecific whose call they received (Greeno & Semple, 2009). Some aspects of chimpanzee sociality are common to male and female chimpanzees—but may vary in their emphasis or relevance between the sexes. Like males, female chimpanzees experience feeding competition—even more so when they are the sole providers to dependent offspring—and might, like subadult and lower ranking males (Clark, 1993), benefit from adjusting their frequency of calling in response to feeding competition.

Similarly, chorusing is associated with social bonding in male chimpanzees (Eckhardt et al., 2015; Fedurek et al., 2014; Mitani & Nishida, 1993; Clark, 1993; Fedurek, Machanda, et al., 2013; Fedurek, Schel, et al., 2013). Given the importance of social bonds to female chimpanzees (Langergraber et al., 2009; Lehmann & Boesch, 2009), their regular use of pant-hoots in chorusing suggests that pant-hoots might serve a similar social bonding function in females. Female chimpanzees may also use pant hoots in ways that are not relevant for males, such as when fostering the vocal and social development of their offspring (Soldati et al., under review). Depending on their offspring’s age or sex, female chimpanzees may alter their own vocal behaviour to avoid conspecifics—and the associated risks of infanticide (Palombit, 2012; Lowe et al., 2020)—or to encourage opportunities for socialisation. For example, in Gombe, female chimpanzees with sons are more gregarious (Murray et al., 2014), potentially providing their male offspring early opportunities to develop the life-long social relationships that impact individual fitness (e.g., Girard-Buttoz et al., 2021). Alternatively, female chimpanzees with maximal oestrus swellings—indicating their period of ovulation—could adjust their likelihood of calling or the acoustic information within their pant-hoots, depending on who is in their immediate party, either to solicit or to avoid potential reproductive partners.

Given the range of sex-specific factors in chimpanzee behaviour (including life history and social differences), the flexibility and range of uses of chimpanzee pant-hoots may be greater than described to date. Identifying the individual and social factors that shape whether and how female chimpanzees produce pant-hoots represents an important step towards a better understanding of the effects of sex-specific behaviour on chimpanzee vocal production. In this study we compare the rate of pant-hoot production in female and male chimpanzees, both when initiating a vocal exchange and when responding to others. We then investigate how individual and socio-ecological factors across sexes and communities mediate the production of pant-hoot responses after hearing the pant-hoots of others.

## Methods

### Ethical statement

Permission to conduct this study was granted by the University of St Andrews Animal Welfare and Ethics Committee, the Budongo Conservation Field Station, the Uganda Wildlife Authority, and the Uganda National Council for Science and Technology (permit number: NS515). We followed the ASAB and IPS guidelines for best practice in field primatology research. This study involved observational data collection only and did not include any direct interaction with the chimpanzees or manipulation of their behaviour beyond our presence.

### Study site

We collected data from Eastern chimpanzees at the Budongo Conservation Field Station (BCFS), in the Budongo Central Forest Reserve, Uganda. This 793km^2^ forest reserve is located at a medium altitude and contains 482km^2^ of continuous and semi-deciduous secondary rainforest cover (Eggeling, 1947). Data were collected from two neighbouring communities of chimpanzees: Sonso and Waibira. At the start of behavioural data collection, the Sonso community consisted of 65 individuals and the Waibira community consisted of 81 named individuals (plus an estimated 20-30 individuals unnamed as very rarely encountered).

### Study subjects

We categorised study subjects as infants (0-4 years old), juveniles (5-9 years old), sub-adults (female: 10-14 years old, male: 10-15 years old), and adults (female: >15 years old, male:>16 years old) following Reynolds (2005). At the beginning of the study, group composition in Sonso was: 34 independent females (27 adult, 7 subadult); 14 independent males (12 adult, 2 subadult); 12 dependent males (3 juvenile, 9 infant); and 13 dependent females (11 juvenile, 2 infant). During the study period, five adults emigrated or died (3 female, 2 male); three subadults matured to adults (2 female, 1 male); eight juveniles matured to subadults (6 female, 2 male); one male infant matured to juvenile; one male infant was born and survived; and three infants were born but died (1 male, 2 unsexed). We collected data from ten independent Sonso females during a total 165 hours of observation (see Supplementary Materials Table S1, S2). The initial group composition in Waibira was: 28 independent females (27 adult, 1 subadult); 26 independent males (18 adult, 8 subadult); ten dependent females (3 juvenile, 7 infant); and 20 dependent males (10 juvenile, 10 infant). During the study period, one juvenile male matured to subadult; four infants matured to juveniles (2 female, 2 male); and five infants were born (2 female, 2 male). We collected data from 20 independent Waibira females during a total 279 hours of observation and from 16 independent Waibira males during a total 171 hours of observation (see Supplementary Materials Table S1, S2).

We considered the potential for sampling bias within our study using the STRANGE framework (Webster & Rutz, 2020). The group size and sex ratio in Sonso are typical for Eastern chimpanzee communities. In Waibira these measures fall at the edge of the Eastern chimpanzee’s range: having both more individuals than typical community and a balanced (rather than female-skewed) sex ratio (c.f., species ranges reported in Wilson et al., 2014). The large group size, which includes a particularly large number of adult males, may influence the ways in which Waibira chimpanzees use their communication to manage social association and any associated resource competition (Eleuteri et al., 2022). In addition, both communities occupy relatively small home ranges, at ∼5.3km^2^ and 10.3km^2^ in Sonso and Waibira respectively (Badihi et al., 2022; Herbinger et al., 2001), which might shape their use of long-distance signals to maintain spatial association across the territory and/or interactions with neighbouring communities.

## Behavioural data collection

### Focal individual follow procedure overview

EH, AS, and CH conducted focal follows with RE, CG conducted focal follows with an experienced field assistant (Altmann, 1974). Focal follows were conducted with 46 individuals between December 2015 to October 2017 between 6am and 6pm (see Supplementary Materials Table S3 for specific dates). Once chimpanzees were located, if there was a choice of more than one adult female to follow, then the focal individual was chosen according to factors prioritised in the following order: how much data were already collected from the individuals present; the oestrus state of females (e.g., oestrus 4 was prioritised as this is the least frequently observed state); their habituation level (with medium to well habituated individuals prioritised); and how commonly they were encountered (with less frequently encountered individuals prioritised). Because the Budongo Central Forest Reserve is a dense secondary rainforest with thick ground cover, continuous observations were challenging and as a result a focal follow could be made up of multiple time periods within the same day (e.g., when the focal was briefly out of sight). A focal follow was defined as data collected from one individual on one day. If data from two individuals were collected in one day, these were assigned separate focal numbers (see statistical methods below).

We collected data on an all-occurrence basis on: focal individual’s activity; party composition changes for focal individual’s party; focal individual pant-hoot production; and pant-hoots heard by the researchers (more details below). As several types of information were being collected on all occurrence basis simultaneously, data were collected by dictating relevant information into a dictaphone supplemented with paper notes. Timings for dictaphone notes were taken from file timestamp information.

### Behavioural context and party composition

We noted information about the behaviour of the focal individual and recorded any change where it lasted 30 seconds or longer. We categorised activity as ‘feeding’ if the focal individual was currently eating, or if they were retrieving or processing food. Activity was categorised as ‘other’ whenever the focal was in a feeding context but not considered to be actively feeding (e.g., they were resting or traveling and not visibly chewing). If the focal individual was feeding, we categorised their food type as either ripe fruit or not ripe fruit (i.e., if eating unripe fruit, leaves, or other foods then food type was categorised as ‘not ripe fruit’).

We recorded the identity and number of all independent individuals in the focal individual’s party continuously (i.e., party composition and size). We considered individuals to be part of the same party as the focal individual if they were within 35 metres of the focal individual (Newton-Fisher, 2004). From this party composition data, we calculated the number of independent individuals present in the party and whether the alpha male of the community was present in the party or not (see ‘Alpha male presence’ below).

### Categorisation of pant-hoots heard and produced by focal individual

The structure of a pant-hoot consists of up to four acoustically distinct phases characteristically described as produced in the following order: introduction, build-up, climax, and let-down (Marler & Hobbett, 1975; Marler & Tenaza, 1977). As individuals often pant-hoot in chorus with others, or after hearing pant-hoots produced by others, pant- hoots can overlap temporally (Fedurek, Machanda, et al., 2013; Marler & Tenaza, 1977), making phases difficult to distinguish, particularly over longer distances. Because these factors might preclude observers from reliably identifying all pant-hoot phases heard, and because recent studies have shown that pant-hoots can be discriminated by the use of any of the phases (Soldati et al., 2022), in this study, we included all pant-hoot vocalisations in which we could discriminate at least one phase of the pant-hoot.

Chimpanzees often produce a pant-hoot after hearing another individual (or several individuals) pant-hoot. A chimpanzee can also produce a ‘spontaneous’ pant-hoot without hearing others pant-hoot. From here on we refer to pant-hooting within one minute after hearing another individual’s pant-hoot, and pant-hooting when joining a chorus that another signaller started, as ‘in response’ pant-hoots (i.e., contrasted with ‘spontaneous’ pant hoots). We only considered pant-hoots produced in response to another pant-hoot as ‘in response’ (not those that followed other vocalizations).

During focal follows, we noted all pant-hoots produced by the focal individual, as well as indicating periods for which we would not have been able to identify whether the focal individual produced a pant-hoot. We also noted the time of all pant-hoots produced by non-focal individuals (in the same party as the focal or in other parties) to: a) note the opportunities focal individuals had to respond to hearing pant-hoots, and b) categorise any pant-hoots that were produced as spontaneous or in response. We operated under the assumption that all calls audible to us were also likely audible to the focal individual, considering that the range of hearing of chimpanzees is comparable to that of humans (Kojima, 1990). As chimpanzees may hear pant-hoots over a greater distance than human observers, the number of pant-hoots we noted as ‘heard’ is likely to be conservative, but equally so across the sex- and community-specific datasets due to the homogenous nature of the landscape across the two territories (e.g., flat terrain, continuous forest).

To understand which features of pant-hoot production elicit pant-hoot responses, we noted: a) whether the pant-hoots heard included at least one pant hoot from within the focal individual’s party or exclusively from outside of it, and b) whether at least one pant- hoot heard was accompanied by drumming. We excluded pant-hoots heard that we estimated to be produced by neighbouring chimpanzee communities on the basis of distance, location, and the chimpanzees’ reaction.

### Opportunities to respond

When a chimpanzee hears a pant-hoot, it gives them an opportunity to respond with a pant- hoot; thus, the number of pant-hoots an individual hears can impact how often they call.

We considered all pant-hoots heard that occurred within less than one minute of the most recent one as part of one event, and thus as one opportunity to respond. Pant-hoots heard after more than one minute without hearing any other pant-hoots were considered a new opportunity to respond. Focal individual pant-hoots were considered as responses if they occurred within one minute of an opportunity to respond. If the focal individual pant- hooted in response, then any pant-hoots heard that started after they finished pant-hooting were considered a new opportunity to respond (even where there was not a one-minute gap since the last non-focal individual pant-hoot heard).

### Female specific traits

Female chimpanzees exhibit sexual swellings throughout their reproductive cycle, with the greatest swelling coinciding with ovulation (Emery & Whitten, 2003). As part of BCFS long-term data collection, experienced field assistants grade the oestrus swellings of female chimpanzees on a scale of 0-4, where zero represents the absence of swelling and four represents the peak of the swelling for that individual. We extracted this information from long-term data for days on which we collected pant-hoot behavioural data from female focal individuals. We then categorised whether a female was in maximal oestrus swelling (grade 4) or not (grades 0 to 3). We also categorised females as parous or nulli-parous (i.e., if they had ever been seen with an infant of their own, or not), and whether they had a dependent male offspring at the time.

### Alpha male presence

To determine the alpha male individual in each community we assessed the dominance hierarchy by calculating Elo-Ratings for each male using the R package ‘EloRating’ (version 0.46.11; Neumann & Kulik, 2020) in R Studio (build 764, ;4.3.2, R Core Team, 2023). Pant-grunts are vocal signals produced by subordinate chimpanzees towards dominant individuals and are widely regarded as a reliable indicator of dominance relationships (Fedurek et al., 2021). We calculated scores from pant-grunts produced by or towards the focal individual that were recorded by field assistants during daily focal follows of male and female chimpanzees between January 2015 and October 2017 (Neumann et al., 2011). Data from 2015 were used as a ‘burn-in’ period to provide more accurate initial rankings at the start of the study period. We calculated the Elo-ratings for each male at the start, middle, and end of the period for which data were used in Models 3 and 4 below. The alpha male was the individual with the highest Elo-rating in all three time periods (see Supplementary Materials Tables S3 – S5).

### Ripe fruit availability FAI methods

Feeding phenology data were extracted from long-term data for the months pant-hoot data were available. Data from two to 30 trees across 18 fruit-bearing feeding species known to be chimpanzee food sources across both territories (total number of trees n = 382), were monitored monthly for ripe fruit. The presence of ripe fruit was scored as available (1) or absent (0). These scores for ripe fruit presence were applied to the following formula to provide monthly ‘Fruit Availability Indices’, or FAI (sensu: Hockings et al., 2010; Takemoto, 2004): FAI = 100 * (∑ (p * f) / ∑ (p * 4)), in which FAI is the fruit availability index percentage, p is the basal area of the tree in cm^2^ and f is the abundance of ripe fruit.

## Statistical analyses

All statistical analyses were run using R Studio (build 764, R version 4.3.2; R Core Team, 2023). Data and code for analyses are available at: https://github.com/Wild-Minds/FemalePantHoots.

We ran four models: Models 1, 3, and 4 were Generalised Linear Mixed Models (GLMMs) fitted using the R package *lme4* (version 1.1-35.1; Bates et al., 2015). Model 2 was a General Linear Model (GLM), fitted using R package *MASS* (version 7.3-54; Venables & Ripley, 2002). Models 1 and 2 were fitted with negative binomial error structure to investigate whether the frequency of pant-hoot production was different for 1) sex or 2) community. Models 3 and 4 were run with binomial error structure to investigate factors that may predict the likelihood of a chimpanzee producing a pant-hoot in response to hearing one or more pant-hoots. Model 3 examined factors which may predict male and female chimpanzee pant-hoot responses. Model 4 examined factors which may predict female chimpanzee pant-hoot responses, including community membership and female-specific variables relating to oestrus status, parity, and the presence of dependent male offspring. A full description of the variables included in each model is described below the general model procedure section.

### General procedure for models

The following general approach was taken for running all models. To improve model interpretation, numeric fixed effects (test and control variables) were z-transformed (i.e., centred around 0 with standard deviation of 1). Non-significant interaction terms (i.e., estimates with *p* > 0.05) were removed from the model so that main effects could be interpreted. We removed random intercepts estimated to be zero to avoid singularity issues. To examine the extent of multicollinearity for predictor variables we calculated the variance inflation factor (VIF) of the fixed effects using the ‘check_collinearity’ function of the package *performance* (version 0.10.8; Lüdecke et al., 2021). Overdispersion was checked for Models 1 and 2 using the ‘check_overdispersion’ function of the package *performance* (version 0.10.8; Lüdecke et al., 2021) (not checked for Models 3 and 4 because the dependent variable for these models was binary). We assessed GLMM stability by comparing the estimates obtained from the model based on all data with those from the model comprising only the fixed control and random effects excluded one at a time using a function generously provided by R. Mundry (Nieuwenhuis et al., 2011). We assessed the overall contribution of fixed effects by comparing each model with a null model comprising only the intercept, control variables, and random effects using a likelihood-ratio test (LRT) (Forstmeier & Schielzeth, 2011). Confidence intervals were derived using the ‘confint’ function of the package *stats* (version 4.3.2; R Core Team, 2023), using 1000 parametric bootstraps and bootstrapping over the fixed and random effects. We used the ‘drop1’ function of the package *stats* (version 4.3.2; R Core Team, 2023) to obtain individual p-values for fixed effects by systematically dropping each fixed effect from the model one at a time and comparing the reduced model with the full model. We omitted test results and p-values of the intercept because of limited interpretation. Because model outputs for interaction effects can be challenging to interpret, where interaction effects were significant (i.e., estimates with *p* > 0.05), we took steps to aid interpretation. Firstly, because the effect on the response of single terms and interaction terms are conditional on the state/value of the other, we visualised these results using the package *interactions* (version 1.1.5; Long, 2019). Secondly, for significant interactions, we conducted post-hoc tests in which we created two data subsets, one for each level of the fixed factor of interest (i.e., ‘sex’ in Model 3) while keeping all other factors from the model the same. For the post-hoc models, we only interpreted main effects for the variables that interacted with the factor of interest in the full model. For overall final models, we determined the marginal (fixed effects) and conditional (fixed and random effects) effect sizes using the ‘model_performance’ function of the package *performance* (version 0.10.8; Lüdecke et al., 2021).

### Models 1 and 2: Are there sex or community differences in the rate of pant-hoot production?

Models 1 and 2 investigated any differences in pant-hoot production frequency for 1) males vs females, and 2) females from Sonso vs females from Waibira, and whether any differences were associated with the number of opportunities to respond. The dependent variable in both models was how many pant-hoots were produced by the focal individual during a focal follow. For both Model 1 and 2, the number of pant-hoots produced was zero inflated, so a negative binomial model error structure was chosen over Poisson. The initial negative binomial models were still over-dispersed when no minimum focal follow length was applied (likely due to severe overdispersion that even the negative binomial structure could not handle). To address this, a minimum focal follow length of 40 minutes was applied. We applied this limit after a visual inspection of a scatter plot of focal follow duration by number pant-hoots produced by focal individual. A minimum focal duration of 40 minutes kept the number of focal follows that had to be excluded to a minimum, while improving the dispersion by excluding short-focal follows with zero pant-hoots produced by the focal individual. When models were run with this minimum focal follow duration, the dependent variable was not over-dispersed (Model 1 dispersion ratio: 1.015; Model 2 dispersion ratio: 1.013).

In Model 1, we tested whether the number of pant-hoots produced was differently predicted for males or females. Sex of the focal individual was included as a fixed factor (reference level: male). To assess whether the number of pant-hoot calls received predicted the number of pant-hoots produced, we also included the number of opportunities to respond as a fixed factor. We initially also included the interaction between the number of opportunities and sex, but this was removed due to non-significance following the procedure described above. As the availability of ripe fruit can impact how much chimpanzees pant-hoot (Notman & Rendall, 2005), and ripe fruit availability varies seasonally, potential pant-hoot production differences between males and females could be explained by variation in food availability across data collection periods. We controlled for food availability in this model by including monthly Food Availability Indices (FAIs) as a control fixed factor. As the amount of time observing an individual will impact how many pant-hoots are observed, we included duration of focal follow (hours) as an offset control factor. When calculating the duration of focal follows, we did not include time when the focal individual was out of sight, or when we would not have been able to identify whether the focal individual produced a pant-hoot. If the focal individual was out of sight for less than ten minutes, we did not consider this as a break in focal length and resumed the focal follow. The identity of the focal individual was included as a random effect. The level of multicollinearity between the independent variables examined was acceptable (maximum VIF value = 1.32). The final structure of Model 1 was as follows:

> Number pant-hoots produced ∼ Sex + Number opportunities to respond + FAI + offset(log(Duration of focal follow)) + (1|ChimpID)

In Model 2, we tested whether the number of pant-hoots produced was predicted differently for Sonso or Waibira females. Community of the focal individual was included as a fixed factor (reference level: Sonso). In this model we also included the number of opportunities to respond as a fixed factor. We also initially included the interaction between the number of opportunities and the community, but this was removed due to non-significance following the procedure described above. We also included monthly FAI as a control fixed factor, and duration of focal follow (hours) as an offset control factor. The model was run as a GLM instead of a GLMM because when identity of the focal individual was included as a random effect it explained negligible variance (<0.001). The level of multicollinearity between the independent variables examined was acceptable (maximum VIF value = 1.28). The final structure of Model 2 was as follows:

> Number pant-hoots produced ∼ Community + Number opportunities to respond + FAI + offset(log(Duration of focal follow))

### Model 3: Which factors predict pant-hoot responses in male and female chimpanzees?

To investigate factors that may predict male and female chimpanzee pant-hoot responses, we fitted a binomial GLMM. This model (Model 3) included data from 28 independent females from the Waibira and Sonso communities and 15 independent males from the Waibira community. Data were not available for Sonso male chimpanzees. The dependent variable included was whether (1) or not (0) the focal individual produced a pant-hoot response after hearing an opportunity to respond (Females: N = 1132 opportunities, 154 responded to; Males: N = 947 opportunities, 159 responded to). The fixed effect variables included in the model were: sex of the focal individual (reference level: male); age of the focal individual (in years); whether the focal was feeding (reference level: no); whether the alpha male was present in the party (reference level: no); the number of independent individuals in the party with the focal individual; whether any of the pant-hoots heard were accompanied by drumming (reference level: no); and whether the pant-hoots heard included one from within the focal individual’s party (reference level: no). We included food type as a nested factor under activity (reference level: not ripe fruit). To investigate which factors varied depending on the sex of the focal individual, we initially included two-way interactions of sex with all other fixed predictor factors. We then removed non-significant interactions one at a time and re-ran the model until only significant interactions remained, so that single effects for variables that were not part of significant interactions could be interpreted. To control for time of day, we included ‘hour’ as a quadratic control factor. As random effects, we included the identity of the focal individual and the number we assigned to the focal follow. Higher order interactions between fixed effects and random slopes were not included due to incomplete combination matrices. Data from the Waibira alpha male were excluded from this model due to our interest in examining whether alpha male presence predicts pant-hoot responses. The median number of independent individuals in the party was four (maximum = 25; however, there were very few instances observed where the number of independent individuals in the party size was 15 or more (37 of 2115 cases); these points were excluded from the model to aid model estimate reliability). The level of multicollinearity between the independent variables examined was acceptable (maximum VIF value: 2.62). The final structure of Model 3 was as follows (where * indicates interactions between variables, and / represents a nested variable)

> Pant-hoot response ∼ Age Centred + Alpha presence + Sex*Pant-hoot received within party + Sex*Independent individuals in party Centred + Drum received + (Activity/Food type) + Hour Centred + I(Hour Centred^2^)+ (1|ChimpID) + (1|FocalFollowNum)

Two post-hoc models were run to further understand the interaction between Sex and Pant-hoot received within party, and Sex and Alpha presence. The first post-hoc model contained only female chimpanzee data, and the second post-hoc model contained only male chimpanzee data. Both post-hoc models were structured as follows:

> Pant-hoot response ∼ Age Centred + Alpha presence + Pant-hoot received within party

> + Independent individuals in party Centred + Drum received + (Activity/Food type) +

> Hour Centred + I(Hour Centred^2^) + (1|ChimpID) + (1|FocalFollowNum)

### Model 4: Which individual factors predict pant-hoot responses in female chimpanzees from two communities?

The fourth model examined factors that may predict female chimpanzee pant-hoot responses, including female-specific variables relating to oestrus status, parity, and the presence of dependent male offspring. We fitted a binomial GLMM that included data from 27 independent females from the Waibira (18 individuals) and Sonso (9 individuals) communities. The dependent variable was whether (1) or not (0) the focal individual produced a pant-hoot response (Sonso: N = 464 opportunities, 52 responded to; Waibira: N =648 opportunities, 92 responded to). As in Model 3, we included the following as fixed factors: age of the focal individual (in years); whether the focal was feeding (reference level: no); whether the alpha male was present in the party (reference level: no); number of independent individuals in the party with the focal individual; whether pant-hoots heard included one from within the focal individual’s party (reference level: no); and whether any of the pant-hoots heard were accompanied by drumming (reference level: no). We also included food type as a nested factor under activity (reference level: not ripe fruit). To control for time of day, we included ‘hour’ as a quadratic control factor. The following additional fixed factors were included in Model 4: the chimpanzee community (reference level: Sonso); whether the female was in maximal oestrus (reference level: no); whether the female was parous (reference level: no); and whether the female had a dependent male offspring (reference level: no). Because only a parous female could have had a dependent male offspring, we included the latter as a nested factor. We looked at two-way interactions of community with each of the other fixed factors. So that single effects for variables not in significant interactions could be interpreted, we removed each non-significant interaction term (estimates with *p* > 0.05) one at a time (starting with the highest p-value) and re-ran the model after each removal until only significant interaction terms remained. We initially included the identity of the focal and the number we assigned to the focal follow as random effects. Focal number was removed from the model due to being estimated as explaining 0 variance. Higher order interactions between fixed effects and random slopes were not included due to incomplete combination matrices. The median number of independent individuals in the party was four (maximum = 25; however, there were few instances observed where the number of independent individuals in the party size was 15 or greater (22 of 1134 cases); and these points were excluded from the model to aid model estimate reliability). The level of multicollinearity between the independent variables examined was acceptable (maximum VIF value: 3.54). The final structure of Model 4 was as follows:

> Pant-hoot response ∼ Age (Centred) + Alpha present + Independent individuals in party (Centred) + Pant-hoot received within party + Drum received + (Activity/Food type) + Maximal Oestrous + (Parity/MaleDep) + Hour Centred + I(Hour Centred^2^) + (1|ChimpID)

## Results

A total of 206 focal follows were conducted on 46 chimpanzees, comprising 616 hours of focal follow data. During focal follows, focal individuals were recorded pant-hooting 527 times and received 2970 pant-hoots (see Supplementary Materials Tables S1 and S2).

Overall, 9.9% (of 232) of pant-hoots produced by females were spontaneous and 30.5% (of 295) pant-hoots produced by males were spontaneous. Mean party size was 5.9 (*SD* = 3.5) independent individuals for Waibira males, 4.7 (*SD* = 3.8) for Waibira females, and 4.4 (*SD* = 3.6) for Sonso females.

### Model 1: Are there sex differences in the rate of pant-hoot production?

After excluding data for individuals that did not meet the minimum criteria for Model 1, we analysed 433 hours of focal data from 118 focal follows on females (29 individuals) and 169 hours from 51 focal follows on males (15 individuals). The full Model 1 explained significantly more variance than the null (LRT: *χ^2^* = 41.1, *df* = 2, *p* < 0.001). The proportion of variance explained by the fixed effects was R^2^marginal = 0.32, and the proportion explained by the fixed and random effects was R^2^conditional = 0.37. Model 1 showed that the number of opportunities to respond predicted how many pant-hoots focal individuals produced (Est = 0.58, *SE* = 0.09, *Z* = 6.50, *p* < 0.001; See Supplementary Materials Table S6 for full model results). Females pant-hooted less frequently (pant-hoots per hour: 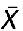 = 0.61, *SD* = 0.58) than males (pant-hoots per hour: 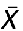 = 1.60, *SD* = 1.13; Model Est = -0.65, *SE* = 0.23, *Z* = -2.87, *p* = 0.007; Figure 1). Food availability did not predict chimpanzees’ pant-hoot production (Est = 0.03, *SE* = 0.11, *Z* = 0.25, *p* = 0.811).

**Figure 1.**
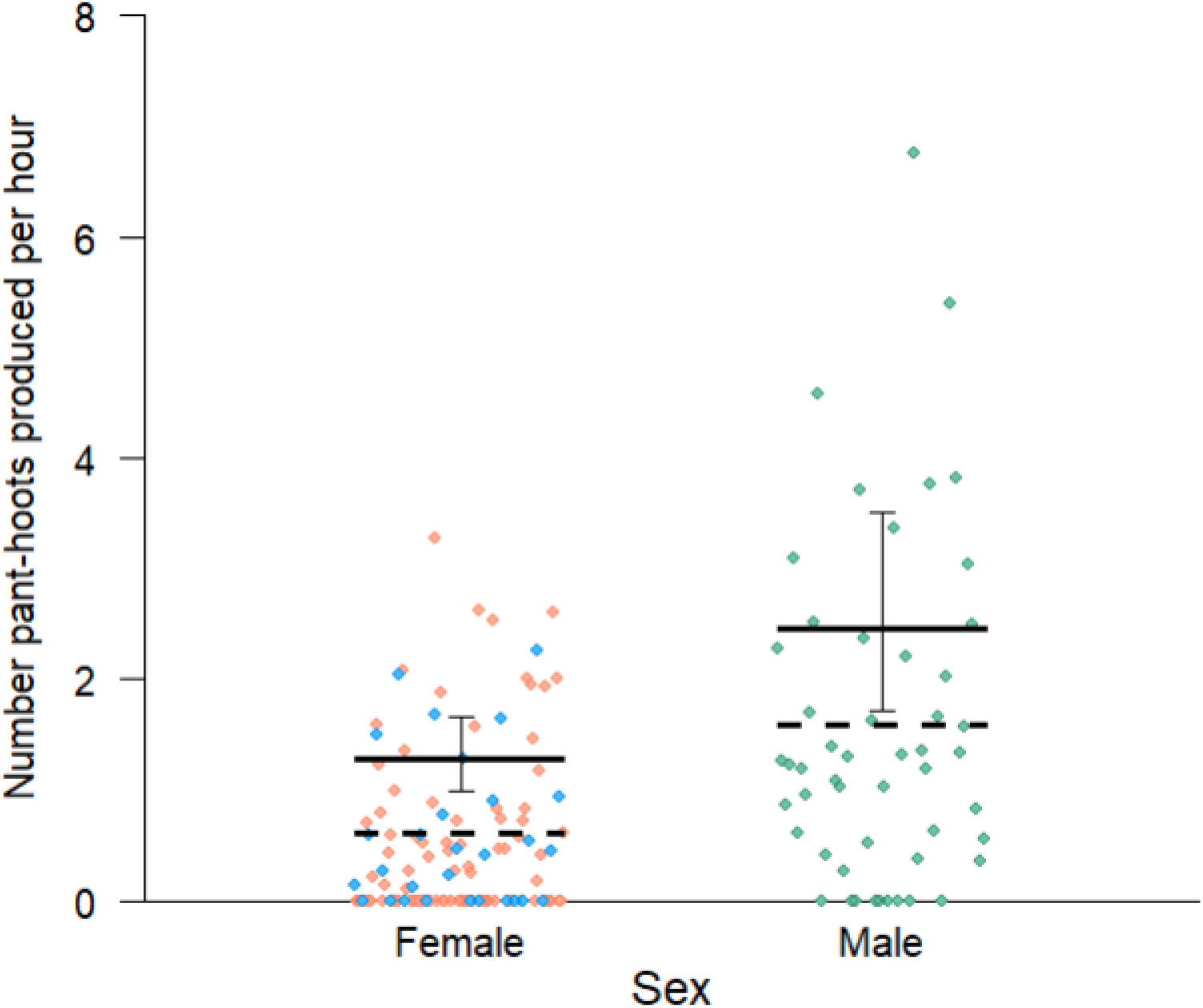
Number pant-hoots produced per hour. Horizontal solid lines represent Model 1 estimates and error bars represent 95% confidence intervals. Dashed lines represent mean values. Dots represent values per focal follow (Nfemale = 118, Nmale = 51). Orange = Waibira females, Blue = Sonso females, Green = Waibira males. Note: while model estimates include contributing factor of ‘number opportunities to respond’, data points and means are not adjusted for the number pant-hoot events received (i.e., they do not express the influence of other factors included in the model).

### Model 2: Are there community differences in female rate of pant-hoot production?

We analysed 268 hours of focal data from 89 focal follows on Waibira females (19 individuals) and 165 hours from 29 focal follows on Sonso females (10 individuals). The full Model 2 explained significantly more variance than the null (LRT *: χ^2^* = 21.3, *df* = 2, *p* < 0.001; Nagelkerke’s *R^2^* = 0.30). The number of opportunities to respond predicted how many pant-hoots focal individuals produced (Est = 0.71, *SE* = 0.13, *Z* = 5.54, *p* < 0.001; See Supplementary Materials Table S7 for full model results). Females in the two communities produced pant-hoots at similar rates (Sonso 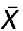 = 0.68, *SD* = 0.60; Waibira 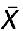 = 0.57, *SD* = 0.59; Model 2 Est = 0.33, *SE* = 0.29, *Z* = 1.13, *p* = 0.263). Food availability did not predict chimpanzees’ pant-hoot production (Est = -0.090, *SE*= 0.14, *Z* = -0.66, *p* = 0.522).

### Model 3: Which factors predict pant-hoot responses in male and female chimpanzees?

We then focused on factors predicting when a focal individual produced pant-hoots in response to hearing pant-hoots. Female chimpanzees (28 individuals) produced pant-hoots in response to 14.1% of 1472 pant-hoot events heard. In the model we excluded data from the alpha male in order to be able to include the variable ‘AlphaPresent’; however, the alpha male responded to 32% of 112 pant-hoot events heard, whereas other male chimpanzees (15 individuals), produced pant-hoots in response to a mean 15.8% of 1058 pant-hoot events heard.

After accounting for data exclusions, 2079 pant-hoot received events were included in Model 3 (N = 1132 for females, N = 947 for males). Model 3 explored factors predicting pant-hoot responding in males and females and explained significantly more variance than the null (LRT: *χ^2^* = 237, *df* = 10, *p* < 0.001). The proportion of variance explained by the fixed effects was R^2^marginal = 0.238, and the proportion explained by fixed and random effects was R^2^conditional = 0.294. We found an interaction between the sex of the focal individual and the number of independent individuals in the party: males were less likely to respond to pant-hoots when they were in larger parties, whereas for females the likelihood of responding was consistent across party size (Table 1; Figure 2A; Tables S8 and S9). We also found an interaction between the sex of the focal individual and whether a pant-hoot was heard from within the party: both males and females were more likely to pant-hoot in response when a pant-hoot was heard from within their party, but this effect was stronger for females (Table 1; Figure 2B; Tables S8 and S9). Tests for outliers within Model 3 indicated there were ten outliers, however when the model was run without outliers, the interpretation of the effects remained the same (see Table S10).

**Figure 2.**
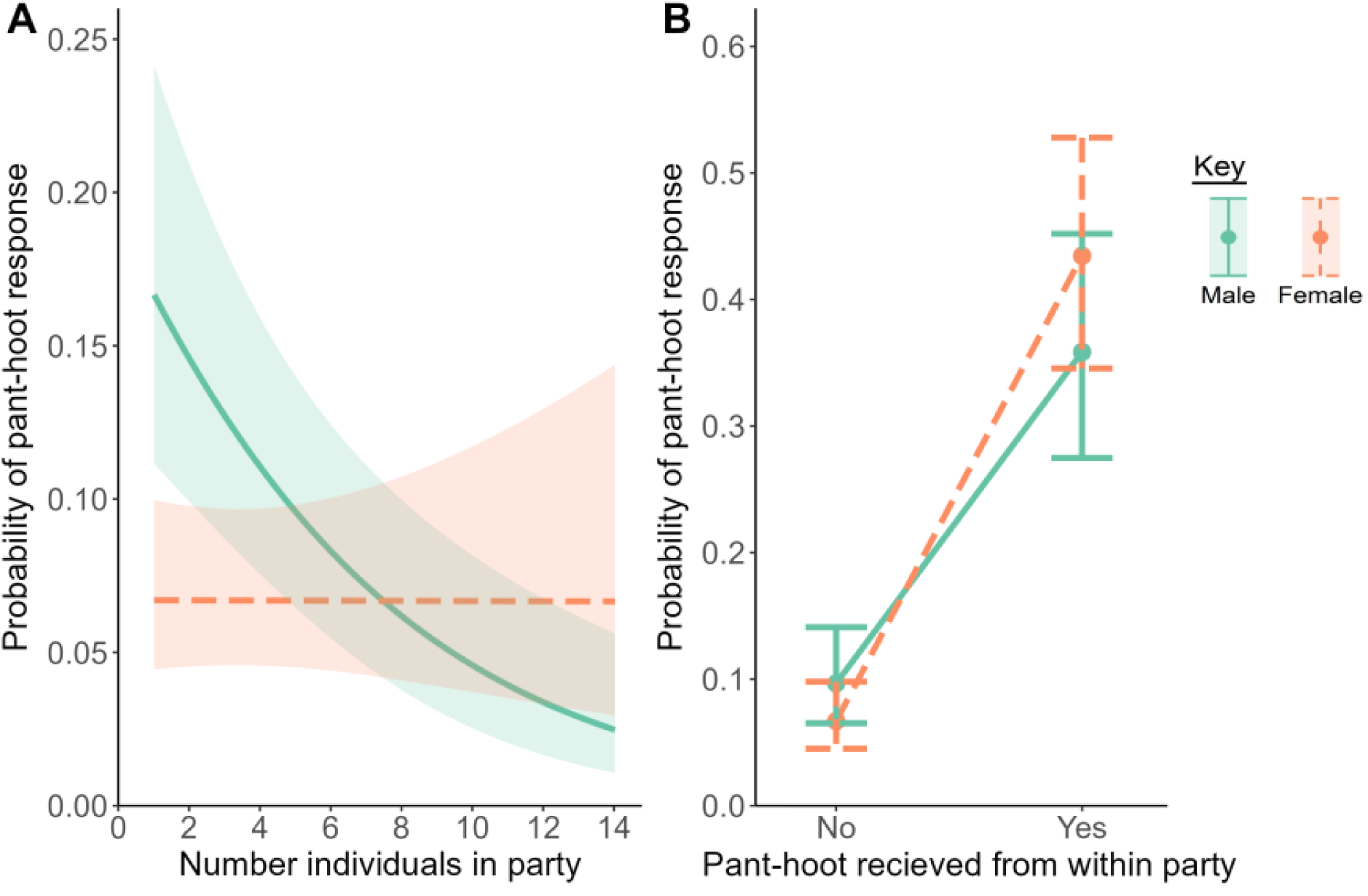
Likelihood of producing a pant-hoot response depending on A) the significant interaction between the focal individual’s sex and number of independent individuals in the party, and B) the significant interaction between the focal individual’s sex and whether a pant-hoot was received from within their party. Error bars and shading represent 95% confidence intervals.

**Table 1.** Parameter estimates for Model 3 exploring factors which predict pant-hoot responses in male and female chimpanzees.

| Model term | Estimate | SE | Z | Lower CI | Upper CI | $\chi^2$ | p |
| --- | --- | --- | --- | --- | --- | --- | --- |
| Intercept | -2.24 | 0.22 | -10.24 | -2.64 | -1.75 |  |  |
| Age (centred) | 0.12 | 0.10 | 1.19 | -0.08 | 0.32 | 1.33 | 0.249 |
| Sex | -0.40 | 0.27 | -1.46 | -0.99 | 0.13 |  |  |
| Alpha present | -0.21 | 0.18 | -1.18 | -0.57 | 0.15 | 1.35 | 0.246 |
| Number individuals in party (centred) | -0.52 | 0.12 | -4.29 | -0.75 | -0.27 |  |  |
| Pant-hoot heard from within party | 1.65 | 0.21 | 7.87 | 1.21 | 2.05 |  |  |
| Drum | -0.07 | 0.17 | -0.40 | -0.40 | 0.27 | 0.16 | 0.686 |
| Activity | -0.75 | 0.26 | -2.92 | -1.26 | -0.17 |  |  |
| Hour of day (centred) | -0.05 | 0.09 | -0.55 | -0.23 | 0.12 | 0.31 | 0.581 |
| l(Hour of day centred)<sup>2</sup> | -0.13 | 0.08 | -1.64 | -0.27 | 0.04 | 2.69 | 0.101 |
| **Sex\*Number individuals in party (centred)** | **0.52** | **0.16** | **3.18** | **0.17** | **0.84** | **10.00** | **0.002** |
| **Sex\* Pant-hoot heard from within party** | **0.72** | **0.30** | **2.36** | **0.14** | **1.30** | **5.54** | **0.019** |
| Activity:Food type | 0.47 | 0.29 | 1.60 | -0.16 | 1.08 | 2.642 | 0.104 |

CI = 95% confidence interval. Interactions are indicated with an asterisk. Nested effects are indicated with a colon. Significant results are depicted in bold.

### Model 4: Which individual factors predict pant-hoot responses in female chimpanzees from two communities?

Sonso females produced pant-hoot responses to 12.2% of pant-hoot events heard (N = 541) and Waibira females produced pant-hoot responses to 15.1% of pant-hoot events heard (N = 931). After accounting for data exclusions, 1112 pant-hoot events received were included in Model 4 (N = 464 for Sonso females, N = 648 for Waibira females). Model 4, which explored factors predicting pant-hoot responding in Waibira and Sonso females, explained significantly more variance than the null (LRT: *χ^2^* = 156, *df* = 11, *p* < 0.001). The proportion of variance explained by the fixed effects was R^2^marginal = 0.30, and the proportion explained by the fixed and random effects was R^2^conditional = 0.30. We found that females were more likely to respond when they heard a pant-hoot from within their party and were less likely to respond when in maximal oestrus (Table 2, Figure 3). Tests for outliers within Model 4 indicated there were two outliers, however when the model was run without outliers, the interpretation of the effects remained the same (see Table S11).

**Table 2.** Parameter estimates for Model 4 exploring factors which predict pant-hoot responses in Waibira and Sonso female chimpanzees.

| Model term | Estimate | SE | Z | Lower CI | Upper CI | $\chi^2$ | p |
| --- | --- | --- | --- | --- | --- | --- | --- |
| Intercept | -1.96 | 0.34 | -5.83 | -2.58 | -1.17 |  |  |
| Community | 0.23 | 0.27 | 0.86 | -0.31 | 0.75 | 0.73 | 0.392 |
| Age (centered) | 0.30 | 0.16 | 1.82 | -0.07 | 0.60 | 3.21 | 0.073 |
| Alpha present | 0.19 | 0.27 | 0.70 | -0.34 | 0.71 | 0.48 | 0.487 |
| Number individuals in party (centered) | 0.01 | 0.11 | 0.13 | -0.19 | 0.25 | 0.02 | 0.898 |
| **Pant-hoot heard from within party** | **2.23** | **0.22** | **10.01** | 1.69 | 2.61 | **111.46** | **< 0.001** |
| Drum | -0.13 | 0.25 | -0.52 | -0.62 | 0.45 | 0.27 | 0.603 |
| Activity | -0.69 | 0.30 | -2.31 | -1.25 | 0.00 |  |  |
| **Maximal oestrous** | **-1.07** | **0.33** | **-3.20** | **-1.65** | **-0.24** | **11.73** | **0.001** |
| Parity | -0.63 | 0.45 | -1.42 | -1.46 | 0.35 |  |  |
| Hour of day (centered) | 0.17 | 0.14 | 1.24 | -0.12 | 0.45 | 1.54 | 0.215 |
| **l(Hour of day** | **-0.30** | **0.11** | **-2.66** | **-0.52** | **-0.03** | **7.15** | **0.007** |

| centered<sup>2</sup>) |  |  |  |  |  |  |  |
| --- | --- | --- | --- | --- | --- | --- | --- |
| Activity: Food type | 0.24 | 0.38 | 0.64 | -0.51 | 0.99 | 0.40 | 0.525 |
| Parity: Male offspring | 0.07 | 0.29 | 0.25 | -0.50 | 0.64 | 0.06 | 0.814 |

CI = 95% confidence interval. Nested effects are indicated with a colon. Significant results are depicted in bold.

## Discussion

We first investigated how the number of pant-hoots produced varied with sex and across females from two communities of wild Eastern chimpanzees. We then investigated how individual and social factors might affect the likelihood of an individual producing pant-hoots in response to hearing the pant-hoots of community members. We found that male and female chimpanzees pant-hooted regularly throughout the day, and heard over five times as many pant-hoots as they produced. Many, but not all, males produced pant-hoots more often than females, highlighting the importance of inter-individual variation as well as overlap between the sexes. Chimpanzees, regardless of their sex, produced pant-hoots more often in response. Excluding the alpha-male, male and female chimpanzees produced in response pant-hoots at similar rates and were three to four times as likely to respond when at least one of the pant-hoots they heard was from within their own party—females especially so. We did not find a difference in the number of spontaneous or response pant-hoots produced by female chimpanzees when comparing neighbouring communities.

Males (but not females) were less likely to produce pant-hoot responses as the number of individuals in their party increased, so much so that while they produced pant-hoots more frequently overall, their responsiveness when in larger parties (>8 individuals) was lower than that of female chimpanzees’. We found no effect of parity, nor the presence of male dependent offspring on female pant-hoot responses, but female chimpanzees were less likely to respond when in maximal oestrus. We found no effect on the probability of pant- hoot responses of the age of the individual, whether alpha male was present in the party, whether the pant-hoots heard were accompanied by a buttress-drum, whether the individual was feeding or not, nor the type of food being consumed.

While, on average, female chimpanzees produce pant-hoots less frequently than males, they do so both spontaneously and regularly in response to hearing pant-hoots as part of intra- and inter-party vocal exchanges. Some female chimpanzees pant-hoot at a higher rate than some males, particularly once opportunities to respond are controlled for. While stating that pant-hoots are not exclusively or predominantly used by males might sound trivial, the exclusion of female pant-hoots from the literature for decades has had important implications for our understanding of chimpanzee long-distance vocal behaviour, with male chimpanzee behaviour used to infer species-typical patterns. Our findings support prior preliminary work suggesting that the use of pant-hoots to take part in vocal exchanges, respond to group members, and potentially coordinate interactions across parties, is not exclusive to male chimpanzees (Clark, 1993; Kalan, 2019; Notman & Rendall, 2005). We show that there are important differences in how social factors predict female, as compared to male, pant-hoot responses. These findings align with more recent perspectives on female chimpanzee behaviour, showing that they are also selective in the way that they navigate social interactions, and regularly participate in activities traditionally associated with male chimpanzees such as hunting (Gilby et al., 2017), border patrols and intercommunity encounters (Samuni et al., 2021), and lethal aggressions (infants: Townsend et al., 2007; adults: Soldati & Hobaiter personal observations). While both males and females use pant-hoots for long- and short-distance communication, the rate at which females pant-hoot seems to be mediated differently by audience effects as well as by female-specific sexual traits. Our findings underline both the importance of studying both sexes to achieve a comprehensive understanding of a species’ communicative behaviour and the mechanisms that shape it, and the risks of introducing bias when generalising species behaviour from only one sex.

The majority of previous pant-hoot studies—and chimpanzee vocal studies more widely— have focused on production, with much less known about receivers and their responses. Given that male pant-hoots have been shown to promote fusion events and recruit other individuals, particularly social partners (during feeding: Bouchard & Zuberbühler, 2022b; travelling: Mitani & Nishida, 1993; and agonistic display: Soldati et al., 2022), females may also respond to pant-hoots in ways that provide information about their location and activity to receivers. Male chimpanzees responded most frequently (and more often than females) in smaller parties. An effect also seen in their long-distance drumming (Eleuteri et al. 2022). They may be using pant-hoots to recruit others—either to increase party size or seeking out a particular composition of individuals (e.g., Fedurek et al., 2014), for example before engaging in higher-risk activities such as patrolling territorial boundaries. In addition, males in smaller parties might particularly benefit from maintaining distant contact with males in other parties, using pant-hoots to coordinate long-distance social dynamics. Male response frequency declined with party size, dropping to very low rates in large parties. It is possible that there is a threshold effect on party-size where the benefits of increased social interactions (Feldblum et al., 2021; Wroblewski et al., 2009) are offset by increased competition—for food or for social partners. Alternatively, given variation in social and hierarchical relationships across individuals, responding may decline in larger parties because the chances of specific preferred partners, or key individuals (such as the alpha male, or a female with oestrus swelling) being already present in the party increases with party size. In contrast, females did not vary their calling with party size—they may similarly have multiple factors at play: larger parties can increase stress (Markham et al., 2014) and feeding competition (Pusey & Schroepfer-Walker, 2013) but also potentially offer greater security.

Both sexes, but females especially so, were more likely to respond when they heard a pant- hoot from within their party, suggesting that they take into consideration their association with nearby, as well as distant, others during vocal exchanges. Responding to the pant- hoots of party members—particularly in choruses, may allow females (opportunities to develop or maintain social relationships with individuals in proximity to them. Calling in chorus, particularly where there is an opportunity for synchronisation, can serve as ‘vocal grooming’ and help mediate long-term social exchanges and relationships with other individuals (Dunbar, 1993; Greeno & Semple, 2009), potentially offsetting the impact of social competition in contexts such as feeding. While there is work showing that spontaneous pant-hoots are predominantly produced when males travel towards, or upon arrival at, a food source (Bouchard & Zuberbühler, 2022; Notman & Rendall, 2005) we found no effect of feeding context. At the same time, as feeding is a stationary activity, individuals who have already pant-hooted at least once may not need to ‘re-advertise’ as neither their location nor activity have changed, especially given that pant-hooting requires that individuals stop feeding to call.

We found that females in maximal oestrous from both communities were less likely to pant-hoot in response than females outwith maximal oestrous, supporting a growing body of evidence in support of flexibility in chimpanzees’ vocal control (e.g., Slocombe et al., 2022; Townsend et al., 2020). Previous studies have shown that female copulation calls do not function to advertise an individual’s oestrus status, but instead inform others about their location and activity, and are produced depending on the identity of individuals in their immediate audience (Fallon et al., 2016; Townsend et al., 2008, 2011). While responding to pant-hoots would be an efficient way for females to broadcast their identity and location to the widest possible range of partners during their fertile period, doing so could also come at a cost. Chimpanzee sexual competition can be intense and includes coercion and aggression by males towards females (Muller & Mitani, 2005). Females in maximal oestrus might strategically refrain from producing pant-hoots to exercise choice over potential reproductive partners by decreasing the likelihood that out-of-sight individuals can identify and locate them across large distances. As males tend to pant-hoot more often in the presence of a female in oestrous (Fedurek et al., 2014), females may be particularly motivated to exert opportunities to control information about their location and status during this critical period.

While reproductive state was predictive of female pant-hoot responses, we did not find an effect of parity, nor of whether mothers had a dependent male offspring at the time.

Research in Gombe chimpanzees suggests that mothers with male offspring spend more time associating and interacting with other group members (Lonsdorf et al., 2014; Murray et al., 2014), and we had hypothesised that females might use pant-hoot responses to facilitate increased social contact when they have male dependent offspring by, for example, recruiting other individuals to their party. However, this was not supported by our data. Recent research suggests higher gregariousness of females with male dependents may not be a universal (Holden, 2021), and this pattern remains untested in the communities we studied. Females with very young infants who are at higher risk of infanticide might pant-hoot less to avoid attracting potentially dangerous males (Lowe et al., 2019), in line with the idea that they tend to isolate themselves around the time of birth (Nishie & Nakamura, 2018). At the same time, mothers might still use others’ pant-hoots to locate males or choose groups to join, without responding with their own calls. Finally, the social benefits of taking part in pant-hoot exchanges could be similarly important for more established parous females as for more recently immigrated nulliparous members of the community. Given the diverse set of factors that appear to influence pant-hoot production, it is challenging to disentangle the rich network of influences to identify the particular immediate mechanisms driving a specific individual to call at any one time. Pant-hoot production likely combines a rich interplay of factors that include competition, risk, safety, or even vocal contagion (Fedurek et al., 2013), as well as social factors.

We found that chimpanzees’ pant-hoot production and responses varied according to their sex. Pant-hoot responses were affected by multiple social factors—including sex-specific ones—indicating that, like those of male chimpanzees, female chimpanzees’ pant-hoots fulfil diverse social functions. Our observations highlight that the same acoustic signal can be used flexibly by both signallers and receivers. Furthermore, our findings showed consistency in the factors that shaped female use of pant-hoots and responses across two communities, despite substantial differences in demography and social structure, suggesting that these effects are largely driven by (sub)species-level features, rather than community- specific cultures. Given their rich acoustic structure, pant-hoots offer substantial opportunities for flexibility in encoding information, exploring the acoustic structure of female chimpanzee pant-hoots will be a key next step.

## Acknowledgments

We are grateful to the management and staff of the Budongo Conservation Field Station (BCFS) for their support on this study. We are especially thankful to the field assistants of the Sonso and Waibira communities, in particular, Geresomu Muhumuza, Monday Mbotella, Chandia Bosco, Adue Sam, Kiiza Vincent, Atayo Gideon, and Mugisha Steven, whose long-term data and expertise contributed to this study. We thank the Royal Zoological Society of Scotland for providing core-funding to BCFS. We thank Roger Mundry for support with the code for statistical analyses. We thank the Uganda Wildlife Authority and the Uganda National Council for Science and Technology for allowing us to conduct this study. The study received funding from The Carnegie Trust for the Universities of Scotland, the European Union’s 8^th^ Framework Programme, Horizon 2020, under grant agreement no802719, and the Kirsten Scott Memorial Trust. We declare no conflict of interest.

