## Supplementary material for "Sex differences in pant-hoot vocalizations in wild Eastern chimpanzees"

### Supplementary materials for: Sex differences in pant-hoot vocalizations in wild Eastern chimpanzees

#### Supplementary methods

##### Study subjects

Table S1 gives an overview of the data collected, including: the number and hours of focal follows, number of individuals, and number of pant-hoots received, and opportunities to respond for females from Waibira and Sonso, and males from Waibira. Table S2 gives a summary for data collected for each individual. Table S3 presents the dates of data collection.

**Table S1.** Summary of data collected for females from Waibira and Sonso, and males from Waibira.

| | N° focal follows | N° focal individuals | N° focal follow hours | N° pant-hoots produced by focal individuals (spontaneous) | N° opportunities to respond |
|---|---|---|---|---|---|
| **Waibira female** | 118 | 20 | 279 | 157 (11) | 1014 |
| **Sonso female** | 29 | 10 | 165 | 81 (12) | 554 |
| **Total female** | 147 | 30 | 444 | 232 (23) | 1568 |
| **Waibira male** | 59 | 16 | 171 | 295 (90) | 1222 |

**Table S2.** Number of focal follows, pant-hoots produced, and opportunities to respond for study subjects.

| Community | Sex | Chimp ID | N° focal follows | N° pant-hoots produced (spontaneous) | N° opportunities to respond |
|---|---|---|---|---|---|
| Sonso | F | AN | 1 | 9 (1) | 49 |
| Sonso | F | IR | 2 | 6 (0) | 11 |
| Sonso | F | KG | 2 | 1 (1) | 18 |
| Sonso | F | KL | 6 | 13 (1) | 119 |
| Sonso | F | KU | 4 | 14 (3) | 64 |
| Sonso | F | KW | 4 | 24 (2) | 123 |
| Sonso | F | KY | 4 | 0 (0) | 43 |
| Sonso | F | NB | 2 | 9 (2) | 20 |
| Sonso | F | OK | 3 | 3 (1) | 54 |
| Sonso | F | WL | 1 | 2 (1) | 53 |
| Waibira | F | AKK | 5 | 14 (1) | 91 |
| Waibira | F | ARU | 3 | 9 (1) | 38 |
| Waibira | F | BAH | 8 | 2 (0) | 13 |
| Waibira | F | JIN | 16 | 17 (1) | 113 |
| Waibira | F | KET | 5 | 8 (0) | 28 |
| Waibira | F | KID | 9 | 8 (1) | 69 |
| Waibira | F | KIP | 4 | 3 (1) | 48 |
| Waibira | F | LIR | 7 | 2 (0) | 34 |
| Waibira | F | LOT | 9 | 20 (5) | 91 |
| Waibira | F | MON | 7 | 12 (0) | 120 |
| Waibira | F | NEV | 5 | 6 (0) | 44 |
| Waibira | F | NOR | 8 | 2 (0) | 48 |
| Waibira | F | ONY | 7 | 3 (0) | 12 |
| Waibira | F | PEN | 4 | 12 (0) | 55 |
| Waibira | F | RAC | 1 | 12 (0) | 37 |

| Community | Sex | Chimp ID | N° focal follows | N° pant-hoots produced (spontaneous) | N° opportunities to respond |
|---|---|---|---|---|---|
| Waibira | F | RIT | 6 | 0 (0) | 20 |
| Waibira | F | SHY | 2 | 3 (1) | 7 |
| Waibira | F | ST2 | 1 | 2 (0) | 48 |
| Waibira | F | TAT | 8 | 22 (0) | 97 |
| Waibira | F | TIB | 3 | 0 (0) | 1 |
| Waibira | M | ALF | 3 | 37 (9) | 100 |
| Waibira | M | ATA | 5 | 21 (6) | 85 |
| Waibira | M | BEN | 3 | 55 (19) | 122 |
| Waibira | M | DOU | 4 | 40 (18) | 96 |
| Waibira | M | FID | 4 | 9 (1) | 78 |
| Waibira | M | GER | 4 | 5 (1) | 68 |
| Waibira | M | KEV | 4 | 6 (0) | 45 |
| Waibira | M | LAF | 4 | 9 (1) | 81 |
| Waibira | M | MAC | 3 | 22 (2) | 89 |
| Waibira | M | MAS | 3 | 9 (1) | 85 |
| Waibira | M | MOR | 4 | 16 (6) | 76 |
| Waibira | M | MUG | 1 | 0 (0) | 3 |
| Waibira | M | SAM | 7 | 6 (1) | 74 |
| Waibira | M | TAL | 3 | 8 (1) | 57 |
| Waibira | M | TRS | 3 | 36 (17) | 100 |
| Waibira | M | URS | 4 | 16 (7) | 63 |

**Table S3** Months of data collection by community. Note date range varies for models due to different inclusion/exclusion criteria.

| Community | Data collection periods | Date range Models 1 & 2 | Date range Models 3 & 4 |
|---|---|---|---|

| | | | |
|---|---|---|---|
| **Waibira** | December 2015 – August 2016<br>December 2016 – May 2017 | 05/12/2015 – 08/10/2017 | 13/01/2016 – 08/10/2017 |
| **Sonso** | February – June 2016 | 16/02/2016 – 04/06/2016 | 16/02/2016 – 04/06/2016 |

#### Dominance hierarchy: Alpha male presence

**Table S4 and S5 give the male dominance hierarchy lists (as measured using Elo ratings) for Waibira and Sonso males for the start, middle, and end of data collection.**

**Table S4** Waibira male dominance hierarchy measured using Elo rating at three time periods. The alpha male is indicated in bold.

| Start | | Middle | | End | |
|---|---|---|---|---|---|
| **ID** | **Elo rating** | **ID** | **Elo rating** | **ID** | **Elo rating** |
| **BEN** | **1** | **BEN** | **1** | **BEN** | **1** |
| URS | 0.784 | URS | 0.918 | KEV | 0.969 |
| ATA | 0.677 | TRS | 0.827 | URS | 0.846 |
| AKL | 0.672 | TAL | 0.767 | TAL | 0.842 |
| TRS | 0.659 | KEV | 0.761 | TRS | 0.738 |
| KEV | 0.586 | AKL | 0.733 | AKL | 0.63 |
| MAP | 0.582 | MAP | 0.566 | MAP | 0.504 |
| CHN | 0.500 | ATA | 0.547 | MAC | 0.496 |
| ALF | 0.491 | CHN | 0.527 | GER | 0.493 |
| GER | 0.409 | ILA | 0.521 | MOR | 0.479 |
| TAL | 0.392 | MOR | 0.506 | DOU | 0.454 |
| KAS | 0.392 | GER | 0.496 | AND | 0.435 |
| MOR | 0.392 | MAC | 0.475 | SDT | 0.435 |
| ABO | 0.392 | ABO | 0.475 | PHI | 0.435 |
| JNO | 0.392 | JNO | 0.475 | MER | 0.435 |
| LIL | 0.392 | LIL | 0.475 | KIL | 0.435 |
| AND | 0.392 | AND | 0.475 | BRT | 0.435 |
| SDT | 0.392 | SDT | 0.475 | LKU | 0.435 |
| PHI | 0.392 | PHI | 0.475 | NOA | 0.435 |
| MER | 0.392 | MER | 0.475 | ROB | 0.435 |
| KIL | 0.392 | KIL | 0.475 | NAL | 0.435 |
| BRT | 0.392 | BRT | 0.475 | MIK | 0.435 |

| Start | | Middle | | End | |
|---|---|---|---|---|---|
| LKU | 0.392 | LKU | 0.475 | BAR | 0.435 |
| NOA | 0.392 | NOA | 0.475 | ALF | 0.427 |
| ROB | 0.392 | ROB | 0.475 | CHN | 0.409 |
| NAL | 0.392 | NAL | 0.475 | JNO | 0.392 |
| MIK | 0.392 | MIK | 0.475 | LIL | 0.385 |
| BAR | 0.392 | BAR | 0.475 | ABO | 0.351 |
| ARD | 0.297 | ALF | 0.463 | ARD | 0.343 |
| DOU | 0.293 | KAS | 0.434 | KAS | 0.301 |
| LAN | 0.284 | DOU | 0.428 | ILA | 0.24 |
| ILA | 0.284 | ARD | 0.34 | DAU | 0.218 |
| SAM | 0.198 | LAN | 0.319 | LAN | 0.186 |
| MUG | 0.194 | MUG | 0.274 | MUG | 0.153 |
| DAU | 0.185 | DAU | 0.21 | SAM | 0.148 |
| LAF | 0.116 | LAF | 0.146 | LAF | 0.119 |
| MAS | 0.013 | SAM | 0.095 | FID | 0.092 |
| FID | 0.009 | MAS | 0.056 | MAS | 0.000 |
| MAC | 0.000 | FID | 0.000 | LAM | NA |
| LAM | NA | LAM | NA | BRI | NA |
| BRI | NA | BRI | NA | | |

Elo rating values were z-standardised.

**Table S5** Sonso male dominance hierarchy measured using Elo rating at three time periods. The alpha male is indicated in bold.

| Start | | Middle | | End | |
|---|---|---|---|---|---|
| **ID** | **Elo rating** | **ID** | **Elo rating** | **ID** | **Elo rating** |
| **HW** | **1** | **HW** | **1** | **HW** | **1** |
| FK | 0.733 | FK | 0.739 | FK | 0.793 |
| MS | 0.691 | MS | 0.699 | MS | 0.738 |
| NK | 0.626 | NK | 0.629 | NK | 0.654 |
| KT | 0.576 | KT | 0.576 | KT | 0.599 |
| SQ | 0.540 | SQ | 0.546 | SQ | 0.565 |
| ZL | 0.479 | ZL | 0.487 | ZL | 0.500 |
| SM | 0.411 | SM | 0.424 | SM | 0.461 |
| MB | 0.335 | MB | 0.335 | ZF | 0.387 |
| ZF | 0.328 | PS | 0.334 | MB | 0.349 |
| PS | 0.299 | ZF | 0.325 | PS | 0.346 |
| JS | 0.291 | JS | 0.224 | JS | 0.229 |
| KZ | 0.214 | KZ | 0.207 | KZ | 0.215 |
| ZG | 0.186 | ZG | 0.186 | ZG | 0.178 |
| KC | 0.157 | KC | 0.157 | KC | 0.164 |
| KS | 0.066 | KS | 0.066 | KS | 0.042 |
| ZD | 0.000 | ZD | 0.000 | ZD | 0.000 |

Elo rating values were z-standardised.

#### Supplementary results

**Model 1: Are there sex differences in the rate of pant-hoot production?**

Table S6 presents the parameter estimates for results of Model 1, a GLMM which explored whether there was difference in the number of pant-hoots produced by male and female chimpanzees when accounting for number of opportunities to respond and Food Availability. A random effect of chimpanzee individual was included.

**Table S6** Parameter estimates for Generalised Linear Mixed Model 1. The outcome variable for this model was number of pant-hoots produced by focal individual within a focal follow, and an oddest factors for duration of focal follow was also included.

| Model term | Estimate | *SE* | *Z* | Lower CI | Upper CI | $X^2$ | *p* |
|---|---|---|---|---|---|---|---|
| **Intercept** | -0.10 | 0.18 | -0.56 | -0.41 | 0.27 | | |
| **Sex** | -0.65 | 0.23 | -2.87 | -1.08 | -0.22 | 7.32 | **0.007** |
| **Number of opportunities to respond** | 0.58 | 0.09 | 6.50 | 0.40 | 0.78 | 33.18 | **<0.001** |
| **FAI** | 0.03 | 0.10 | 0.25 | -0.18 | 0.24 | 0.06 | 0.811 |

CI = 95% confidence interval. Significant results are depicted in bold.

**Model 2: Are there community differences in the rate of female pant-hoot production?**

Table S7 presents the parameter estimates for results of Model 2, a GLM which explored whether there was difference in the number of pant-hoots produced by female chimpanzees from Waibira and Sonso communities when accounting for number of opportunities to respond and Food Availability.

**Table S7** Parameter estimates for General Linear Model 2. The outcome variable for this model was number of pant-hoots produced by focal individual within a focal follow, and an oddest factors for duration of focal follow was also included.

| Model term | Estimate | *SE* | *Z* | Lower CI | Upper CI | $X^2$ | *p* |
|---|---|---|---|---|---|---|---|
| **Intercept** | -1.12 | 0.25 | -4.45 | -1.61 | -0.61 | | |

| | | | | | | | |
|---|---|---|---|---|---|---|---|
| **Community** | 0.33 | 0.29 | 1.13 | -0.25 | 0.90 | 1.25 | 0.263 |
| **Number of opportunities to respond** | 0.71 | 0.13 | 5.54 | 0.41 | 1.03 | 23.6 | **<0.001** |
| **FAI** | -0.09 | 0.14 | -0.66 | -0.37 | 0.19 | 0.41 | 0.522 |

CI = 95% confidence interval. Significant results are depicted in bold.

**Model 3: Which factors predict pant-hoot responses in male and female chimpanzees?**

Table S8 and S9 show the parameter estimates for Model 3 post-hoc tests, where Table S8 model only included data from female focal individuals, and Table S9 model only included data from male focal individuals. These post-hoc models were run to further understand the interaction between a) Sex and Number individuals in party, and b) Sex and pant-hoot received from within the party. These results show that number of independent individuals in the party did note predict female pant-hooting in response, but showed that that males were less likely to pant-hoot in response when they are in larger parties. These results also showed that both females and males more likely to pant-hoot in response if they receive a call from within their party, but that this effect was stronger for females than males. A random effect of chimpanzee individual, and focal follow number was included.

**Table S8** Parameter estimates for post-hoc test of the interactions in Model 3 using only data from female individuals. The outcome variable for this model was whether a pant-hoot was produced by the focal individual after hearing an opportunity to respond.

| **Model term** | **Estimate** | ***SE*** | ***Z*** | **Lower CI** | **Upper CI** | ***$X^2$*** | ***p*** |
|---|---|---|---|---|---|---|---|
| Intercept | -2.55 | 0.21 | -11.86 | -2.94 | -2.08 | | |
| Age (centered) | 0.09 | 0.12 | 0.80 | -0.13 | 0.34 | 0.62 | 0.431 |
| Alpha present | 0.13 | 0.27 | 0.47 | -0.40 | 0.67 | 0.25 | 0.618 |
| **Number individuals in party (centered)** | **-0.05** | **0.12** | **-0.38** | -0.26 | 0.19 | **0.28** | **0.595** |
| **Pant-hoot heard from within party** | **2.36** | **0.22** | **10.54** | 1.89 | 2.76 | **124.84** | **<0.001** |
| Drum | -0.31 | 0.26 | -1.22 | -0.80 | 0.23 | 1.53 | 0.217 |
| Activity | -0.73 | 0.30 | -2.39 | -1.30 | -0.03 | | |
| Hour of day (centered) | 0.14 | 0.13 | 1.06 | -0.14 | 0.41 | 1.10 | 0.294 |
| I(Hour of day centered)$^2$ | -0.22 | 0.10 | -2.23 | -0.40 | 0.01 | 5.19 | 0.023 |

| | | | | | | | |
|---|---|---|---|---|---|---|---|
| Activity:Food type | 0.49 | 0.37 | 1.32 | -0.33 | 1.20 | 1.91 | 0.167 |

CI = 95% confidence interval. Interactions are indicated with an asterisk. Nested effects are indicated with a colon. Significant results are depicted in bold.

**Table S9** Parameter estimates for post-hoc test of the interactions in Model 3 using only data from male individuals. The outcome variable for this model was whether a pant-hoot was produced by the focal individual after hearing an opportunity to respond.

| **Model term** | **Estimate** | ***SE*** | ***Z*** | **Lower CI** | **Upper CI** | ***$X^2$*** | ***p*** |
|---|---|---|---|---|---|---|---|
| Intercept | -2.33 | 0.26 | -8.88 | -2.80 | -1.75 | | |
| Age (centered) | 0.25 | 0.17 | 1.46 | -0.11 | 0.59 | 2.01 | 0.156 |
| Alpha present | -0.52 | 0.24 | -2.17 | -1.00 | -0.01 | 4.65 | 0.031 |
| **Number individuals in party (centered)** | **-0.44** | **0.13** | **-3.40** | -0.69 | -0.18 | **12.17** | **<0.001** |
| **Pant-hoot heard from within party** | **1.71** | **0.21** | **8.04** | 1.25 | 2.12 | **72.55** | **<0.001** |
| Drum | 0.15 | 0.23 | 0.67 | -0.29 | 0.60 | 0.43 | 0.511 |
| Activity | -0.86 | 0.49 | -1.76 | -1.71 | 0.73 | | |
| Hour of day (centered) | -0.25 | 0.13 | -1.88 | -0.52 | 0.01 | 3.47 | 0.062 |
| I(Hour of day centered)$^2$ | -0.04 | 0.12 | -0.34 | -0.27 | 0.22 | 0.11 | 0.737 |
| Activity:Food type | 0.71 | 0.52 | 1.35 | -1.00 | 1.67 | 1.98 | 0.160 |

CI = 95% confidence interval. Interactions are indicated with an asterisk. Nested effects are indicated with a colon. Significant results are depicted in bold.

**Table S10.** Parameter estimates for version of Model 3 run when outliers were excluded.

| **Model term** | **Estimate** | ***SE*** | ***Z*** | **Lower CI** | **Upper CI** | ***$X^2$*** | ***p*** |
|---|---|---|---|---|---|---|---|
| Intercept | -2.29 | 0.24 | -9.65 | -2.72 | -1.73 | | |
| Age (centered) | 0.15 | 0.11 | 1.41 | -0.06 | 0.37 | 1.86 | 0.172 |
| Sex | -0.60 | 0.31 | -1.95 | -1.23 | 0.00 | | |
| Alpha present | -0.28 | 0.19 | -1.53 | -0.64 | 0.11 | 2.28 | 0.131 |
| Number individuals in party (centered) | -0.58 | 0.13 | -4.64 | -0.82 | -0.28 | | |
| Pant-hoot heard from within party | 1.76 | 0.21 | 8.20 | 1.29 | 2.18 | | |
| Drum | -0.13 | 0.18 | -0.75 | -0.48 | 0.24 | 0.56 | 0.455 |
| Activity | -0.77 | 0.27 | -2.86 | -1.24 | -0.20 | | |
| Hour of day (centered) | -0.08 | 0.10 | -0.85 | -0.28 | 0.10 | 0.70 | 0.402 |
| I(Hour of day centered)$^2$ | -0.11 | 0.08 | -1.43 | -0.26 | 0.06 | 2.03 | 0.154 |
| **Sex*Number individuals in party (centered)** | **0.54** | **0.17** | **3.14** | 0.15 | 0.87 | **9.72** | **0.002** |
| **Sex* Pant-hoot heard from within party** | **0.90** | **0.32** | **2.83** | 0.23 | 1.47 | **8.00** | **0.005** |

| | | | | | | | |
|---|---|---|---|---|---|---|---|
| Activity:Food type | 0.33 | 0.31 | 1.08 | -0.32 | 0.91 | 1.18 | 0.277 |

CI= 95% confidence interval. Interactions are indicated with an asterisk. Nested effects are indicated with a colon. Results for variables which were part of a significant interaction with Sex in the full Model 3 are depicted in bold.

#### Model 4: Which individual factors predict pant-hoot responses in female chimpanzees from two communities?

**Table S11.** Parameter estimates for version of Model 4 run when outliers were excluded

| Model term | Estimate | *SE* | *Z* | Lower CI | Upper CI | $X^2$ | *p* |
|---|---|---|---|---|---|---|---|
| Intercept | -1.90 | 0.34 | -5.61 | -1.90 | -1.68 | | |
| Community | 0.25 | 0.28 | 0.89 | 0.00 | 0.49 | 0.78 | 0.377 |
| Age (centered) | 0.33 | 0.17 | 2.01 | 0.05 | 0.50 | 3.92 | 0.048 |
| Alpha present | 0.15 | 0.27 | 0.54 | -0.04 | 0.29 | 0.29 | 0.592 |
| Number individuals in party (centered) | -0.01 | 0.11 | -0.05 | -0.01 | -0.01 | 0.00 | 0.960 |
| **Pant-hoot heard from within party** | **2.31** | **0.23** | **10.19** | **2.19** | **2.48** | **117.01** | **< 0.001** |
| Drum | -0.18 | 0.26 | -0.71 | -0.39 | 0.00 | 0.51 | 0.475 |
| Activity | -0.66 | 0.30 | -2.20 | -0.96 | -0.55 | | |
| **Maximal oestrus** | **-1.12** | **0.34** | **-3.27** | **-1.46** | **0.00** | **12.36** | **< 0.001** |
| Parity | -0.82 | 0.46 | -1.80 | -0.98 | -0.81 | | |
| Hour of day (centered) | 0.17 | 0.14 | 1.21 | 0.00 | 0.34 | 1.45 | 0.229 |
| **I(Hour of day centered)$^2$** | **-0.31** | **0.12** | **-2.70** | **-0.40** | **-0.23** | **7.47** | **0.006** |
| Activity:Food type | 0.06 | 0.39 | 0.16 | 0.00 | 0.12 | 0.02 | 0.875 |
| Parity:Male dependent offspring | 0.20 | 0.30 | 0.65 | 0.00 | 0.39 | 0.38 | 0.538 |

CI = 95% confidence interval. Interactions are indicated with an asterisk. Nested effects are indicated with a colon. Significant results are depicted in bold.
